# Systematically attenuating DNA targeting enables CRISPR-driven editing in bacteria

**DOI:** 10.1101/2022.09.14.507927

**Authors:** Daphne Collias, Elena Vialetto, Jiaqi Yu, Khoa Co, Éva d.H. Almási, Ann-Sophie Rüttiger, Tatjana Achmedov, Till Strowig, Chase L. Beisel

**Author notes:** Correspondence (to C.L.B.).

## Abstract

Bacterial genome editing commonly relies on chromosomal cleavage with Cas nucleases to counter-select against unedited cells. However, editing normally requires efficient recombination and high transformation efficiencies, which are unavailable in most strains. Here, we show that systematically attenuating DNA targeting activity enables RecA-mediated repair in different bacteria, allowing chromosomal cleavage to drive genome editing. Attenuation can be achieved by altering the format or expression strength of guide (g)RNAs; using nucleases with reduced cleavage activity; or engineering attenuated gRNAs (atgRNAs) with disruptive hairpins, perturbed nuclease scaffolds, non-canonical PAMs, or guide mismatches. These modifications greatly increase cell counts and even improve the efficiency of different types of edits for Cas9 and Cas12a in *Escherichia coli* and *Klebsiella oxytoca*. We further applied atgRNAs to restore ampicillin sensitivity in *Klebsiella pneumoniae*, establishing a new resistance marker for genetic studies. Attenuating DNA targeting thus offers a counterintuitive means to achieve CRISPR-driven editing across bacteria.

## INTRODUCTION

The study and engineering of bacteria have been vastly improved with CRISPR technologies and the ongoing advances in CRISPR-based editing^1,2^. Traditionally, bacteriophage-derived DNA recombinases are combined with RNA-guided Cas nucleases to achieve efficient editing ranging from base changes to large deletions and insertions^3,4^. The recombinases drive the homologous recombination of a DNA repair template (RT), while the Cas nucleases counterselect against unedited cells through the generation of cytotoxic double-stranded DNA breaks to the unedited bacterial chromosome^3^. The flexibility of this approach allows for both small and large edits and thus remains the method of choice for editing in bacteria despite a growing set of other options^5– 9^. However, CRISPR-based counterselection typically requires high transformation efficiencies and the availability of compatible phage-based recombinases unavailable in most bacteria. Even with improvements that delay counterselection with inducible DNA targeting^10–13^ or limit bacterial escape with RecA inhibitors^14^, CRISPR-based counterselection still remains largely off-limits outside of model bacteria and, even in model bacteria, incredibly difficult for challenging edits such as larger insertions, multiplexed editing, and libraries.

Despite the paradigm of utilizing chromosomal cleavage to counterselect against unedited cells, prior work reported an intriguing exception: repair of chromosomal cleavage by Cas9 in *Escherichia coli* through endogenous homologous recombination^15^. Recombination was hypothesized to come from weaker targeting at certain sites, which left uncleaved copies of the chromosome that could mediate homologous recombination and maintain cell viability. Recombination was dependent on RecA and, in one instance, drove integration of a plasmid-encoded RT in the absence of a heterologous recombinase. Under this setup, driving homologous recombination of a supplied RT would achieve flexible editing while circumventing the need for heterologous recombinases or high transformation efficiencies due to enhanced cell survival. However, survival was seemingly random and site-dependent, creating uncertainties about whether CRISPR-driven homologous recombination could be broadly achieved in bacteria.

Here, we show that CRISPR-driven homologous recombination can be systematically achieved by attenuating DNA targeting activity, in different cases boosting the number of transformants as well as the editing efficiency. The most tunable approach, which involved what we call attenuated gRNAs (atgRNAs), could achieve flexible editing with Cas9 and Cas12a nucleases in different bacteria and mediate small base exchanges and large insertions. The approach obviated the need for a heterologous recombinase and greatly increased the number of recovered colonies without sacrificing editing efficiency, the two major drawbacks of traditional recombination-based editing in bacteria. The use of atgRNAs thus represents a new paradigm for CRISPR-based editing in bacteria that achieves improved editing outcomes by scaling back targeting activity.

## RESULTS

### Altering gRNA format and expression can boost cell counts and genome editing

We were initially intrigued why targeting some locations in the *E. coli* genome with the *Streptococcus pyogenes* Cas9 led to RecA-dependent homologous recombination rather than cell death^15^. One observation from this work was that the tested gRNAs were all encoded as CRISPR arrays, with each array expressed from its native promoter from *S. pyogenes*. The transcribed arrays are processed into CRISPR RNAs (crRNAs) through the action of a separate trans-activating crRNA (tracrRNA)^16^. In contrast to the CRISPR arrays, single-guide RNAs (sgRNAs), which circumvent the need for a separate tracrRNA, are commonly used for Cas9-based editing in bacteria. Each sgRNA is also normally expressed from a heterologous promoter^11,14,17,18^. We refer to the processed crRNA:tracrRNA duplex and the sgRNA as gRNAs. We thus asked if this difference in gRNA format or expression accounts for RecA-mediated cell survival, helping us work toward CRISPR-driven editing in bacteria.

Using a standard plasmid transformation assay in *E. coli* in which a gRNA plasmid is transformed into cells already harboring a Cas9 plasmid (**Fig. 1A**), we targeted genomic sites that previously yielded different extents of cell survival in the presence of *recA*^*15*^. Under this setup, cell counts of the WT and *recA* strains depend on targeting activity: cells unable to undergo repair would yield low cell counts in the presence or absence of *recA*, cells undergoing RecA-mediated repair would yield high colony counts in the presence of *recA* and low colony counts in the absence of *recA*, and cells with limited targeting would yield high colony counts even in the absence of *recA*. Similar to the prior work, applying the transformation assay using minimal CRISPR arrays driven by the native promoter (Pn RSR) yielded varying extents of survival in the presence of *recA* and consistently low survival in the absence of *recA* compared to a non-targeting (NT) control (**Fig. 1B**). In contrast though, sgRNAs driven by a constitutive synthetic promoter (Ps sgRNA) consistently yielded virtually no cell counts in the presence or absence of *recA* for the same genomic target sites (**Figs. 1B and S1A**). Target locations alone therefore cannot explain RecA-mediated cell survival and instead point to factors related to gRNA format and expression.

**Figure 1:**
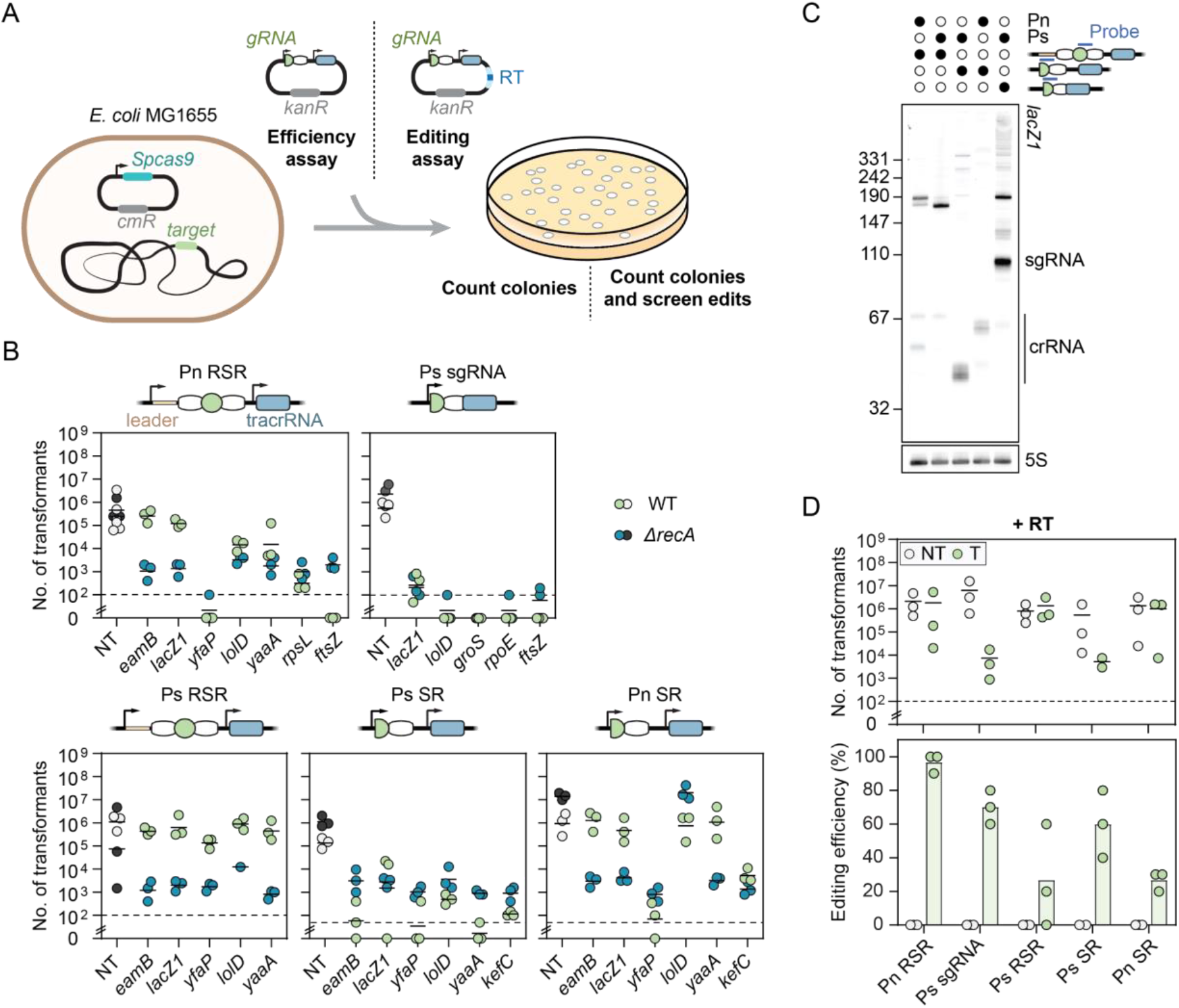
The outcome of chromosomal targeting in *E. coli* depends on gRNA format and abundance. **A**) Schematic of the experimental setup for chromosomal targeting and editing. **B**) Genome targeting assay in *E. coli* and *E. coli* Δ*recA*. **C**) Northern blot analysis of whole RNA isolated from *E. coli* Δ*lacI-Z* with pCas9 and pgRNA. A *lacZ1* spacer specific probe was used to probe the abundance of each RNA product. A 5S probe was used as a control. The approximate size of an sgRNA is indicated to the left of the Northern blot and the approximate size of mature crRNAs are indicated with a line. **D**) Genome editing assay in *E. coli* targeting *lacZ1* to introduce AvrII restriction enzyme recognition site as a silent mutation. Individual dots for the transformations indicate a single biological replicate. Dashed lines indicate the limit of detection from plating. NT and T indicate targeting and non-targeting gRNAs, respectively. Individual dots for the editing efficiencies indicate the average of 3 colonies screened from one biological replicate for NT samples or 10 colonies screened from one biological replicate for targeting samples.

To tease apart gRNA format and expression, we replaced the native promoter of the CRISPR array with the constitutive synthetic promoter (Ps RSR) (**Fig. 1B**). We also expressed a semi-processed crRNA with either promoter (Ps SR, Pn SR) to interrogate any contributions from processing. Combining the synthetic promoter with the native array (Ps RSR) or the native promoter with the semi-processed crRNA (Pn SR) generally yielded elevated colony counts in the presence of *recA*, while combining the synthetic promoter with the semi-processed crRNA (Ps SR) yielded low colony counts in the presence of *recA*. Colony counts were consistently low in the absence of *recA*, underscoring the importance of RecA-mediated homologous recombination. These data suggest that changing the format and expression of the gRNA can alter the outcome of survival.

The operating hypothesis is that weaker targeting allows cells to survive through RecA-mediated recombination with replicated copies of the genome^15^. Following this hypothesis and the observed impact of colony counts on gRNA format, we would expect weaker targeting to derive from lower gRNA abundance that could direct fewer Cas9 molecules to cut target DNA, allowing RecA-mediated repair to outcompete extensive DNA cutting. In line with our expectation, Northern blotting analysis on the complete set of gRNAs targeting *lacZ1* revealed an inverse correlation between the abundance of the final gRNA (Ps sgRNA > Ps SR > Pn SRS ≈ Pn SR > Ps RSR) with colony counts in the presence of *recA* (**Fig. 1C**). The crRNA sizes also varied, possibly contributing to the extent of DNA targeting. Similar trends in gRNA size and abundance were observed targeting two other locations in *lacZ* (**Fig. S1B-D**). These results provide direct evidence that reduced gRNA abundance can lead to cell survival through RecA-mediated recombination in the absence of a provided RT.

If the cells survive genome targeting by Cas9 via RecA-mediated recombination, then the presence of a RT whose edit prevents targeting should lead to genome editing without sacrificing colony counts. We therefore introduced a plasmid-encoded RT to mutate part of the *lacZ1* target into a restriction site and assessed colony counts and the editing efficiency (**Figs. 1D and S1E**). All gRNA formats yielded variable colony counts that paralleled those in the absence of the RT, with Ps sgRNA and Ps SR yielding the fewest colonies and the other formats (Pn RSR, Ps RSR, Pn SR) yielding the most colonies. Intriguingly, Pn RSR yielded the highest editing efficiency (∼97%) even above that from Ps sgRNA (70%), while the other formats yielded more variable yet measurable editing (**Fig. 1D**). In total, changing gRNA format and expression can achieve CRISPR-driven editing that boosts colony counts, obviates the need for an exogenous recombinase, and can preserve the efficiency of precise editing.

### Systematically attenuating genome targeting with Cas9 can increase colony counts and editing efficiencies

The impact of gRNA format and expression suggested that purposefully attenuating DNA targeting activity could not only increase colony counts but also even improve the editing efficiency. If the goal is to attenuate targeting, we can envision multiple means that weaken or delay any step between gRNA and Cas9 production to DNA cleavage (**Fig. 2A**), including altering the expression or stability of the gRNA or Cas nuclease, slowing nuclease:gRNA complex formation, and reducing DNA target recognition or cleavage. Furthermore, we reasoned that any approach could be implemented individually or together to fine-tune the activity reduction. We therefore tested various attenuation approaches beyond modifying gRNA format to determine if any of these approaches could predictably boost cell counts and editing.

**Figure 2:**
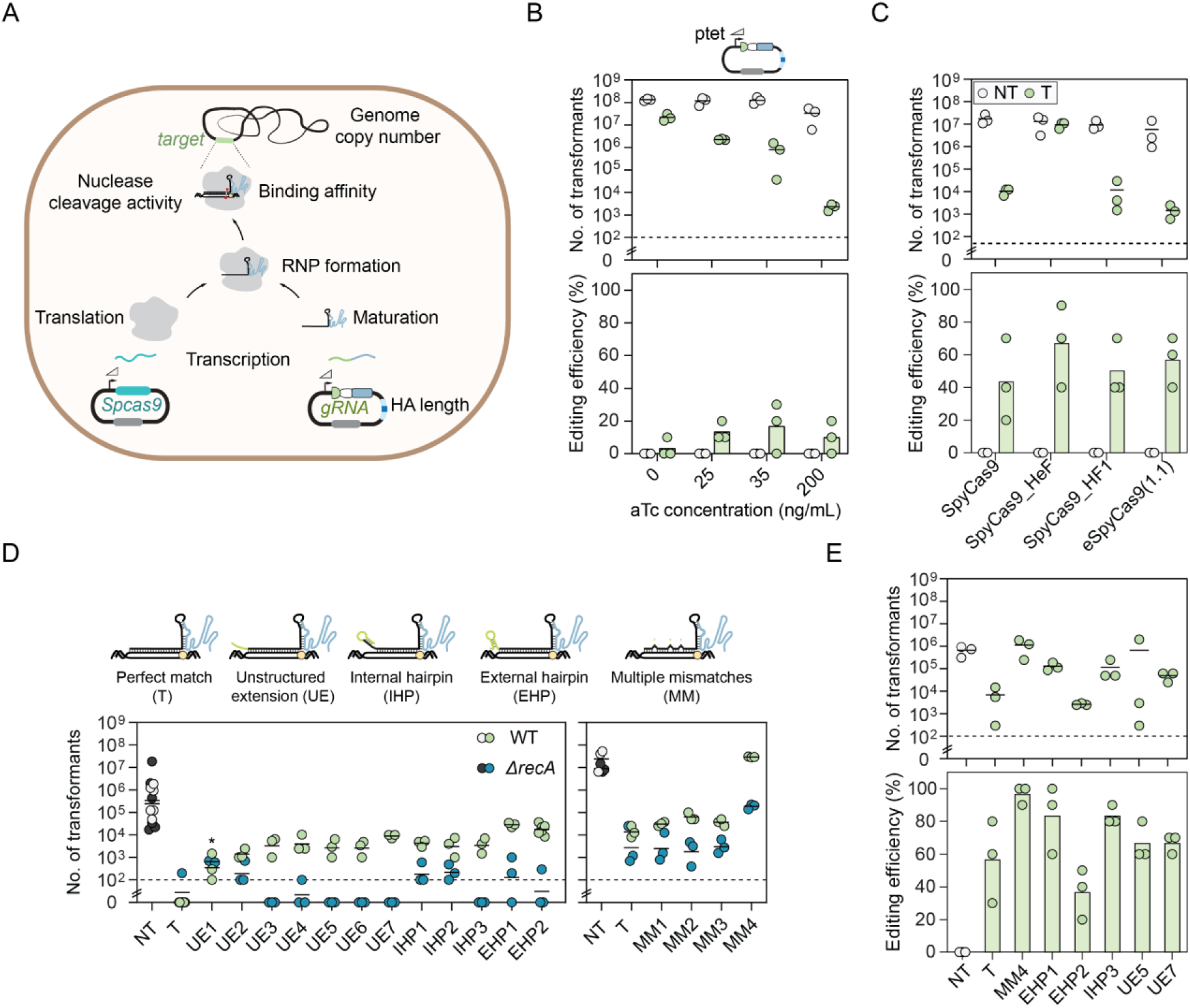
Modulating DNA targeting activity, including through attenuated gRNAs, can boost colony counts without sacrificing editing efficiencies. **A)** Schematic of different steps that can be altered to attenuate DNA targeting to improve homologous recombination with a supplied RT. The maturation step applies only to crRNAs. **B)** Genome editing assay in *E. coli* with aTc inducible sgRNA expression. A non-selective out-growth with aTc induction was used prior to selective plating. **C)** Genome editing assay in *E. coli* using WT, HeF, HF1, and e(1.1) SpyCas9. **D)** Genome targeting assay in *E. coli* and *E. coli* Δ*recA* with modified sgRNAs and Cas9. **E)** Genome editing assay in *E. coli* using selected attenuated gRNAs (atgRNAs) from D). Cells were made electrocompetent at ABS_600_ ≈ 1.6 for the editing assay. Individual dots for the transformations indicate a single biological replicate. Dashed lines indicate the limit of detection from plating. NT indicates non-targeting and T indicates targeting. Individual dots for the editing efficiencies indicate the average of 3 colonies screened from one biological replicate for NT samples or 10 colonies screened from one biological replicate for targeting samples. * indicates that the transformants resulted in a lawn or uncountable colonies.

We first tested the impact of modulating sgRNA expression using a tetracycline-inducible promoter (**Fig. 2B**). This approach offered excellent flexibility, as different concentrations of the inducer anhydrotetracycline (aTc) could be readily added to tune targeting activity. Accordingly, titrating the aTc concentration in cultures transformed with the sgRNA, Cas9, and RT plasmids resulted in colony count reductions varying between ∼6-fold (0 ng/ml aTc) and ∼10^5^-fold (200 ng/ml aTc) compared to the NT control. When introducing mutations into the *lacZ1* target to create a restriction site, consistent editing was observed even with lower aTc concentrations without sacrificing colony counts. However, the extent of editing never exceeded an average of 20% even for the highest applied aTc concentration.

Next, we tested the impact of reducing DNA targeting activity with high-fidelity Cas9 nucleases (**Fig. 2C**). These nucleases were evolved to more readily reject mismatches and therefore reduce off-target DNA cleavage^19–21^. However, they also tended to exhibit lower cleavage efficiencies at otherwise perfect targets, which we hypothesized could promote RecA-mediated survival upon genome targeting even with an sgRNA conferring robust DNA targeting. Testing three established high-fidelity versions of Cas9 (SpyCas9_HeF, SpyCas9_HF1, eSpyCas9(1.1)) along with a constitutive synthetic sgRNA that resulted in extensive cell killing (**Fig. S1B**), we found that SpyCas9_HeF exhibited much higher colony counts than Cas9 and the other two variants (∼1,000-fold), yet all variants exhibited similar average editing efficiencies (∼50-60%) compared to Cas9 (∼43%). Less-active nucleases, such as some high-fidelity nucleases, therefore offer a distinct means of achieving CRISPR-driven editing, although some nucleases outperform others.

As a final approach, we reasoned that modifying the gRNA itself to reduce Cas9 binding, target recognition, or target cleavage could be applied to achieve CRISPR-driven editing. Fortunately, numerous modifications are known that can have a minor to massive impact on DNA targeting, including introducing mismatches into the guide sequence at different locations (SM or MM)^22^, extending the PAM-distal end of the gRNA to include an unstructured extension (UE) or a hairpin that does (IHP) or does not (EHP) extend into the guide sequence^23,24^, targeting genomic sequences adjacent to non-canonical PAMs (NC)^3,25–28^, and mutating the fixed region of the gRNA bound by Cas9 to affect RNA folding or Cas9 recognition^29^ (**Figs. 2D and S1F**). Using the *lacZ3*-targeting sgRNA as the starting point, the modifications gave varying degrees of colony counts in the presence or absence of *recA*. For most of the modifications though, at least one variant yielded elevated colony counts in the presence versus absence of *recA*, indicating attenuated activity. Combining a subset of these modified variants with the RT to introduce a restriction site in *lacZ* led to editing efficiencies similar or even higher to the original sgRNA but with modestly to greatly increased colony counts (**Fig. 2E**). For instance, one approach in which three target mismatches were introduced into the sgRNA guide (MM4) yielded colony counts indistinguishable from a non-targeting sgRNA along with ∼97% editing. In total, sufficiently attenuating DNA targeting by Cas9 through different means can increase colony counts while preserving the editing efficiency via RecA-mediated homologous recombination.

### CRISPR-driven editing with atgRNAs is generalizable beyond Cas9

While Cas9 is arguably the most widely used nuclease for genome editing in bacteria, a growing suite of DNA-targeting Cas nucleases are becoming available^30^ whose targeting activity could be attenuated to enhance genome editing in bacteria. One increasingly popular DNA-targeting nuclease is Cas12a^31^. Apart from being structurally and functionally distinct from Cas9, Cas12a can process a transcribed CRISPR array without any accessory factors^31^ and recognizes T-rich PAMs^31^ distinct from those recognized by Cas9^32^. Cas12a was also employed in one bacterial application when Cas9 proved cytotoxic^33^. We therefore asked if atgRNAs specific to Cas12a could be generated to achieve CRISPR-driven editing in bacteria.

We focused on the widely-used Cas12a from *Acidaminococcus* species (AsCas12a)^31^ given its widespread use for CRISPR technologies. After confirming that designed gRNAs consistently lead to low cell counts when targeting the *E. coli* genome in the absence of *recA* (**Fig. S2A**), we generated variants of a *lacZ*-targeting gRNA in line with the Cas9 atgRNAs: appending an external hairpin on the PAM-distal end of the guide (EHP), introducing a mutation in the Cas12a repeat shown to partially inhibit DNA targeting^34^ (mutR), mutating the leader region upstream of the Cas12a repeat to introduce a hairpin that can also inhibit DNA targeting^35^ (IRH), targeting sites with non-canonical PAMs (NC)^32^, and introducing a single target mismatch (SM) or multiple target mismatches (MM)^22^ in the guide sequence (**Figs. 3A and S2B-C**). As with Cas9, at least one variant for most of the tested modifications yielded elevated colony counts in the presence versus absence of *recA*. Using two of these variants (**Fig. 3B**), we assessed genome editing by introducing either a silent mutation that generates a restriction site or a two-nucleotide deletion at the *lacZ* target site (**Figs. 3C and S2D-E**). In both cases, the atgRNAs yielded ∼1,000-fold greater colony counts compared to the original gRNAs. Critically, the editing efficiencies were also much higher for the atgRNAs (95-100%) versus the original gRNAs (10-30%). High colony counts and editing was lost in the absence of *recA* (**Fig. S2F**), confirming the importance of RecA-mediated homologous recombination with atgRNAs. Therefore, CRISPR-driven editing in bacteria can be achieved with atgRNAs that extend beyond Cas9.

**Figure 3:**
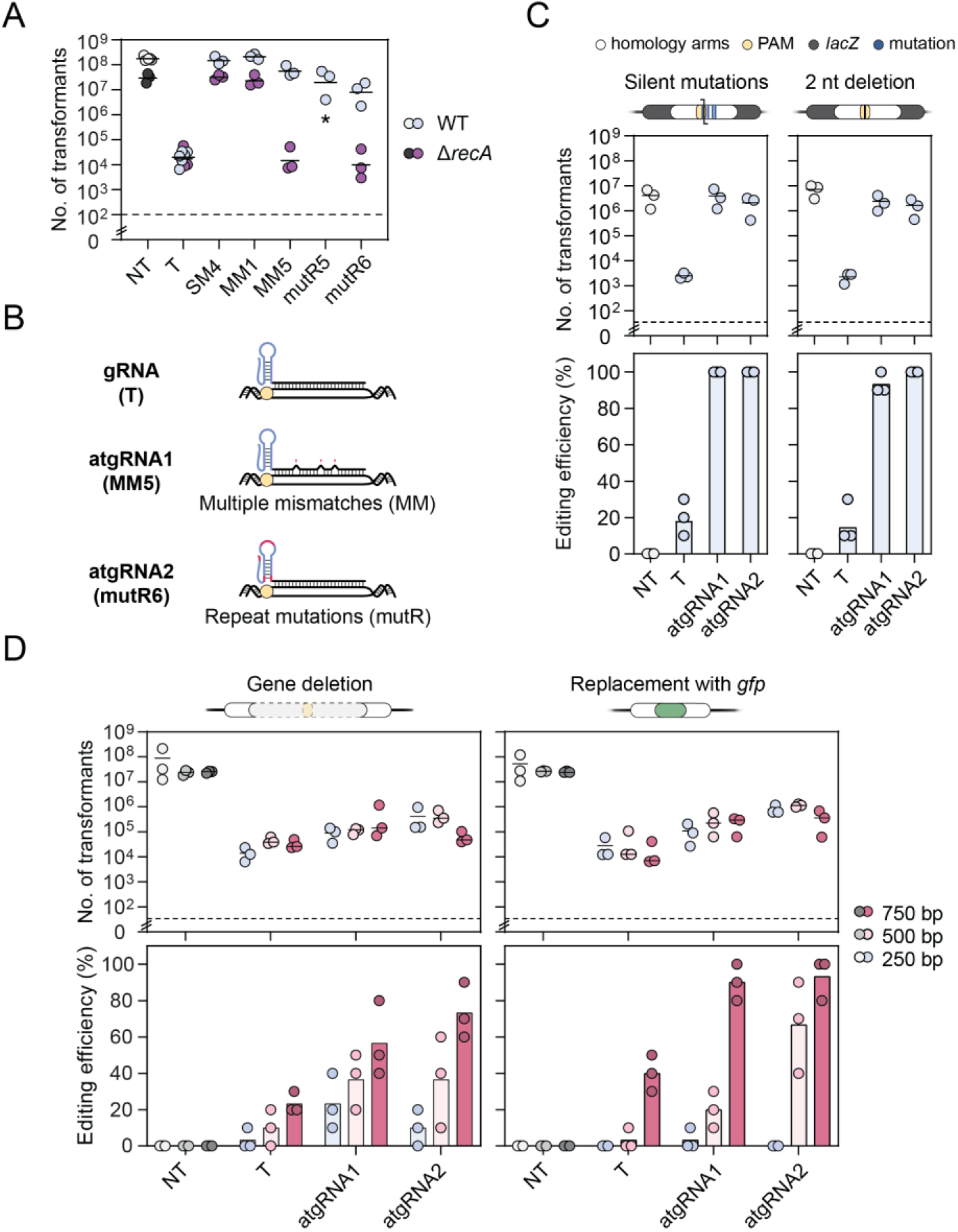
CRISPR-driven editing with attenuated gRNAs extends to Cas12a. **A)** Genome targeting assay in *E. coli* and *E. coli* Δ*recA* with modified gRNAs and Cas12a. **B)** Schematic of atgRNAs for Cas12a. **C)** Genome editing assay in *E. coli* using atgRNAs to introduce a RE silent mutation (left panels) and a 2nt deletion in the *lacZ* ORF (right panels). **D)** Genome editing assay in *E. coli* using atgRNAs to delete the entire *lacZ* gene (left panels) and substitute *lacZ* with *gfp* (right panels). Individual dots for the transformations indicate a single biological replicate. Dashed lines indicate the limit of detection from plating. NT indicates non-targeting and T indicates targeting. Individual dots for the editing efficiencies indicate the average of 3 colonies screened from one biological replicate for NT samples or 10 colonies screened from one biological replicate for targeting samples. * indicates that the transformants resulted in a lawn or uncountable colonies.

Given the superior performance of Cas12a atgRNAs over gRNAs when generating small edits, we asked how they perform when generating larger edits. We attempted two different types of large edits: the deletion of *lacZ* (∼ 3 kb) and the replacement of *lacZ* with a 717-bp fragment encoding *gfp* (**Figs. 3D and S2G**). As these types of edits are typically more difficult to create, we varied the length of the homology arms of the RT to help improve the efficiency of homologous recombination^36–38^. While the atgRNAs did not fully recover colony counts for these large edits, they still yielded higher colony counts than the original gRNA (e.g., 13-fold for atgRNA1 and 20-fold for atgRNA2 for *gfp* replacement with 750-bp homology arms). The atgRNAs also always yielded higher editing efficiencies than the original gRNA, with longer homology arms consistently increasing the editing efficiency. Therefore, when testing more challenging edits, atgRNAs can outperform the original gRNA in both colony counts and editing efficiencies.

### atgRNAs can improve editing performance in *Klebsiella* species

Building on the demonstrations of CRISPR-driven editing in *E. coli*, we asked how well this approach could extend to bacteria with less developed tools. As a first demonstration, we focused on *Klebsiella oxytoca*, a commensal bacterial species recently shown to reduce the intestinal colonization of multidrug-resistant (MDR) *Klebsiella pneumoniae* in various mouse models^39^. While Cas9-based editing with the lambda-red recombination system had been implemented in one strain^40^, this approach required extensive sgRNA screening and generally resulted in few transformants. Applying atgRNAs therefore offered an opportunity to expand the editing tools in this clinically relevant bacterium while also setting the stage to further interrogate the mechanistic basis of colonization resistance.

As the test case within the *K. oxytoca* strain MK01, we sought to delete a 2.7-kb fragment of the *cydAB* operon previously shown to be important for growth under microaerophilic conditions in *E. coli* ^*41*^. We began with the established Cas9 editing system^40^ to provide a direct basis of comparison and because the absence of lambda-red did not yield any detectable edits with a linear or plasmid RT (**Figs. 4A and S3A-B**). While the tested four sgRNAs yielded different extents of colony counts, one sgRNA (sp47) yielded the lowest colony counts and no detectable edits (**Fig. 4B**). Circumventing the more extensive modification testing we performed in *E. coli*, we opted to introduce the hairpin EHP2 to the PAM-distal end of each sgRNA and assessed editing (**Fig. 4B and S3D**). For the sp47 sgRNA, the atgRNA boosted colony counts to be consistently above the limit of detection and yielded an editing frequency of ∼83% (**Fig. 4B**). For the other sgRNAs, either addition of the external hairpin maintained colony counts and high editing frequencies (sp53) or greatly boosted the colony counts but at the expense of editing (sp51, sp55). In these cases, DNA targeting was likely too weak to drive editing, at least within the timeframe of the experiment. Therefore, expedient testing of one atgRNA modification enhanced editing, although more extensive screening of atgRNA modifications could further enhance editing across target sites.

**Figure 4:**
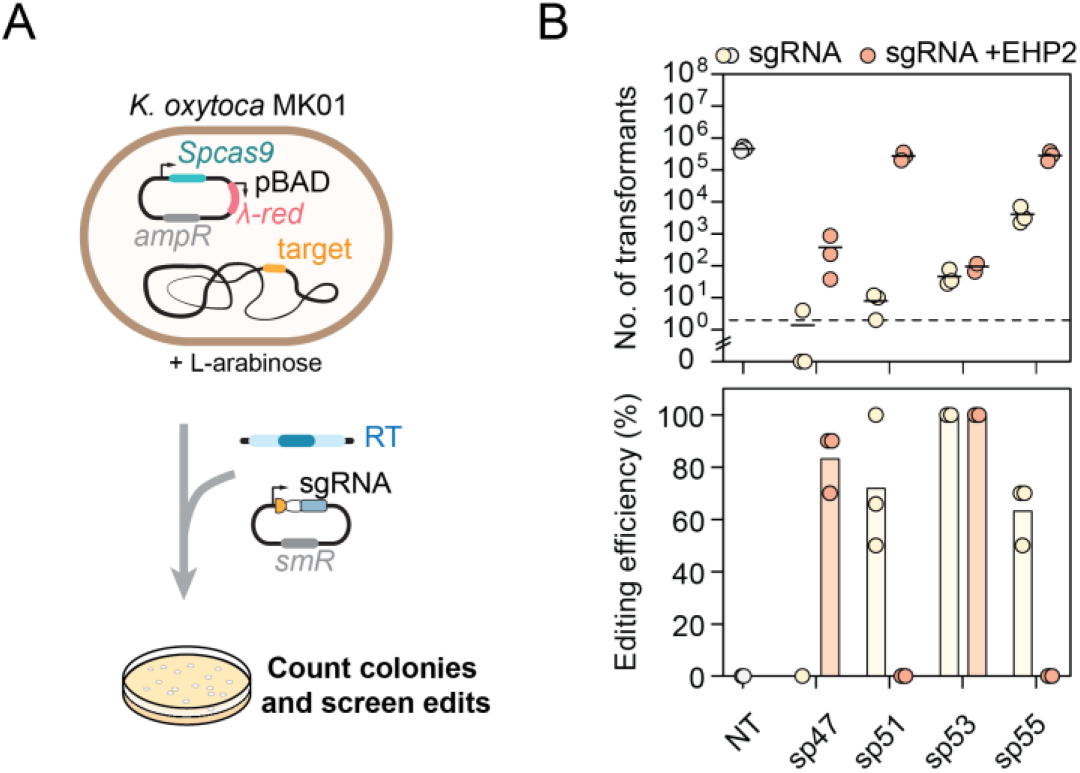
An attenuated gRNA enhances editing in *Klebsiella oxytoca*. **A)** Schematic of genome editing experimental setup. **B)** Genome targeting assay in *K. oxytoca* with sgRNAs and atgRNAs to delete *cydAB*. Individual dots for the transformations indicate a single biological replicate. Dashed lines indicate the limit of detection from plating. NT indicates non-targeting. Individual dots for the editing efficiencies indicate the average of 3 colonies screened from one biological replicate for NT samples or 10 colonies screened from one biological replicate for targeting samples.

Beyond *K. oxytoca*, we turned to a second demonstration of CRISPR-driven editing outside of *E. coli*: eliminating an antibiotic resistance marker in *Klebsiella pneumoniae* to facilitate new genetic tools. *K. pneumoniae* is commonly associated with antibiotic resistance, complicating treatment of infections. At the same time, resistance greatly restricts which antibiotics can be applied to select for plasmids, limiting genetic studies and efforts to combat future infections.

We sought to reverse resistance to ampicillin by disrupting *blaSHV-1*, an intrinsic β-lactamase gene common to *K. pneumoniae* strains^42^. We focused on the commonly studied strain ATCC 10031 that is sensitive to chloramphenicol but not ampicillin (**Fig. 5A**). A Cas12a gRNA targeting *blaSHV-1* yielded few colonies in the presence or absence of *recA*, offering a starting point to generate atgRNAs (**Fig. S3E**). Of the tested atgRNAs, two harboring target mismatches in the guide and one containing a mutated Cas12a repeat were combined with a mutation in the constitutive gRNA promoter led to elevated colony counts in the presence versus the absence of *recA* (**Fig. S3E**). Proceeding with these three atgRNAs and an RT intended to generate a premature stop codon (**Fig. 5B**), we found that all three yielded >1,000-fold increase in colony counts compared to the original gRNA (**Fig. 5C**). In addition, one of the atgRNAs (kpn_atgRNA3) yielded ∼95% editing compared to ∼10% editing with the original gRNA. The other two atgRNAs yielded similarly low editing as the original gRNA, albeit with much greater colony counts that facilitates finding an edited colony.

**Figure 5:**
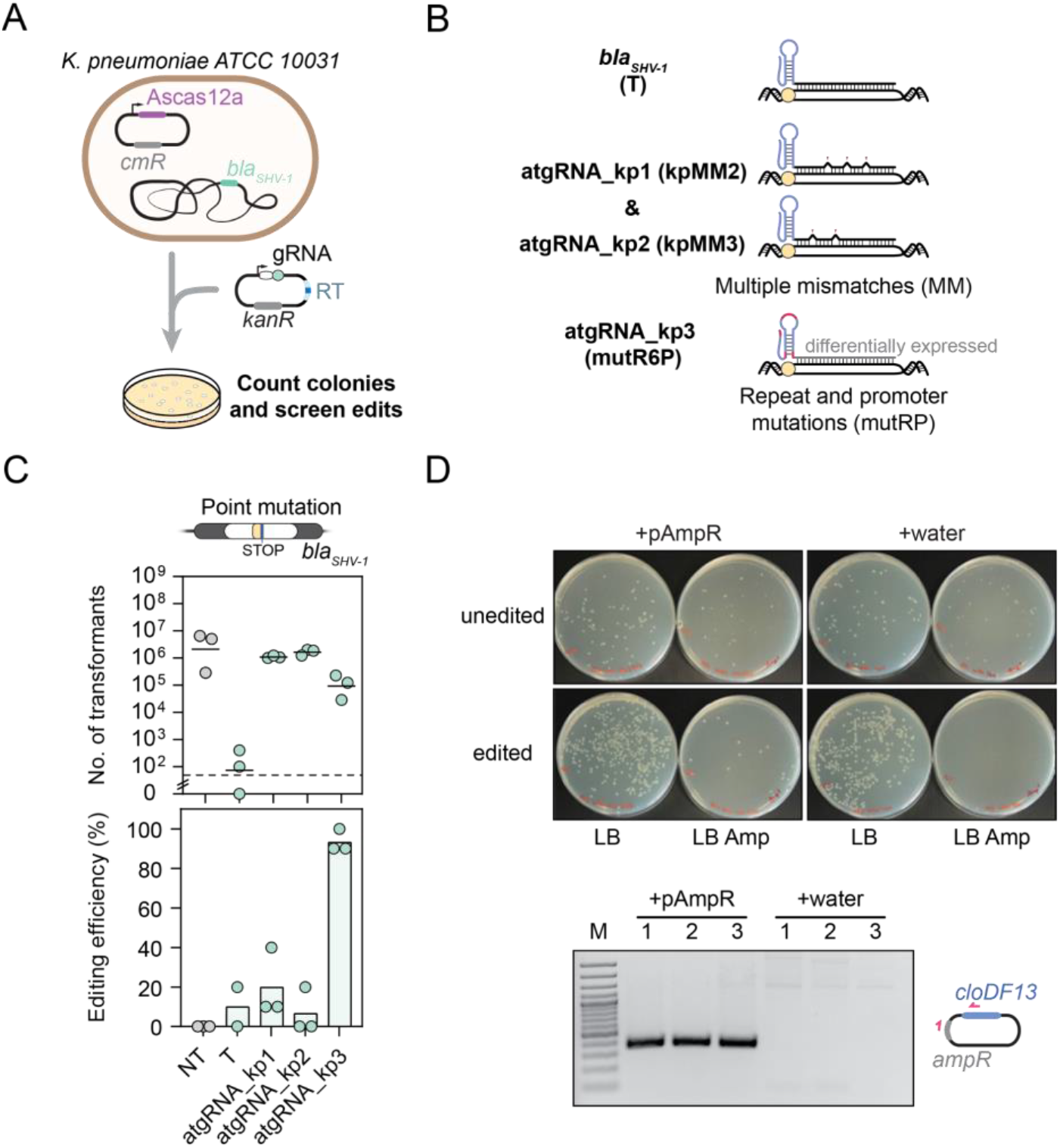
Attenuated gRNAs reverse antibiotic resistance in multidrug-resistant *Klebsiella pneumoniae*. **A)** Schematic of genome editing assay. **B)** Schematic of canonical and attenuated guide design. **C)** Genome editing assay in *K. pneumoniae* to introduce a stop codon in the *bla*_SHV-1_ gene. Each dot in the editing plot is the result obtained after screening 10 and 3 colonies respectively for the T and NT samples. For the T gRNA, each dot represents the editing efficiency obtained for 4 colonies due to the absence of other colonies on the plates. One of the biological replicates yielded no colonies for this condition. **D)** Top: plate images upon transformation of edited cells (atgRNA_kp3) and unedited cells (NT) with the pAmpR plasmid or water in Amp or LB only plates. Bottom: schematic of pAmpR amplification by PCR with confirmatory gel image. The three lanes on the left are edited cells transformed with pAmpR, while the three lanes on the right are edited cells transformed with water (from LB plates). M: marker.

Continuing with the strain edited with atgRNA3, we evaluated transformation of the pAmpR plasmid conferring resistance to ampicillin (**Fig. 5D**). The edited strain yielded colonies only in the presence of pAmpR, while the wildtype strain yielded colonies in the presence or absence of pAmpR when grown on plates containing ampicillin. Screening edited colonies that formed with ampicillin confirmed the presence of pAmpR. The edit therefore expands the genetic toolbox available to this strain with an additional selectable marker. Overall, we conclude that atgRNAs can be applied to achieve CRISPR-driven editing in non-model bacteria even for large edits, in some cases outperforming the more traditional CRISPR-based counterselection.

## DISCUSSION

In this work, we showed that systematically attenuating DNA targeting activity can achieve CRISPR-driven editing in bacteria, greatly boosting colony counts and even increasing the frequency of precise genome editing. This seemingly paradoxical concept--making DNA targeting weaker can improve editing--appears to emerge from the cells having time to repair cut genomic DNA with other copies of the chromosome or a provided RT. Repair through genomic DNA is likely more efficient, although repair with the RT prevents retargeting and thus drives the eventual build-up of edited cells in the population. Because of cell survival and the large number of resulting transformants, CRISPR-driven editing could enable the generation of edits normally challenging if not off-limits in bacteria, such as the generation of large libraries or simultaneous multiplexed edits. In contrast, efficient DNA targeting leads to rapid cutting of all genomic copies, forcing cells to undergo efficient homologous recombination immediately or eliminating cells that did not undergo recombination. In either case, efficient DNA targeting would also select for escape mutants in the population that are resistant to CRISPR targeting (e.g., mutated Cas9 plasmid or gRNA plasmid)^43–45^, accounting for the sometimes large fraction of unedited cells with a standard gRNA.

Achieving CRISPR-based editing in bacteria draws helpful parallels to editing in mammalian cells. There, DNA cuts engage different DNA repair pathways, allowing CRISPR to drive editing. In mammalian cells though, repair is dominated by end-joining pathways that lead to seemingly random edits. When relying on these pathways (e.g., for gene disruption), editing has proven to be highly efficient and can be readily multiplexed to disrupt numerous genes in one pass^46^. Achieving precise edits through homologous recombination in mammalian cells however requires either extensive screening or a series of approaches to enhance this repair pathway or inhibit other repair pathways^47–50^. As bacteria broadly lack the necessary machinery for end joining^51^, homologous recombination instead represents the dominant repair pathway. For this reason, attenuating DNA targeting to enhance editing is likely unique to bacteria, despite the parallels to CRISPR-driven editing in human cells.

One challenge with attenuating DNA targeting in bacteria is finding the “sweet spot” that sufficiently drives RecA-mediated homologous recombination. If targeting is too strong, then a large population of cells will be killed off; if targeting is too weak, then few of the cells will undergo editing. Our work and prior work suggest that the extent of attenuation needed to hit this sweet spot likely varies between gRNA:target site pairs as well as organisms^15^. Fortunately, a large set of options are available to attenuate DNA targeting as illustrated in this work, with atgRNAs posing the most flexible and fine-tuned means. To find the most appropriate option, we recommend first testing representative atgRNAs (e.g., target mismatches, repeat mutations, gRNA format) in the WT and *recA*-deletion strain to identify modifications that greatly boost colony counts in the presence of *recA*; if a *recA* deletion cannot be obtained in the strain-of-interest, a dominant-negative RecA or homologous recombination inhibitor (e.g., GamS) can be co-expressed to inhibit the endogenous repair pathways^14,52^. Furthermore, we reason that, upon transformation with the atgRNA + RT, further culturing should continuously boost editing efficiencies by giving cells more opportunities to repair DSBs using the RT. In the future, we envision high-throughput screens combined with machine learning to predict the best atgRNA (or set of atgRNAs) for a given site and desired edit, paralleling work developing efficient guides for different applications^53–55^. Given the utility of atgRNAs in *E. coli* and in *Klebsiella* species, we anticipate that these approaches could be broadly applied across the bacterial world, facilitating future mechanistic studies and engineering efforts.

## METHODS

### Plasmid and strain construction

Table S1 contains all strains, plasmids, and oligonucleotides used in this work. The *E. coli* MG1655 Δ*recA* (*recA*) and MG1655 Δ*lacI-Z* (*lacI-Z*) strains were produced using Flp-FRT recombination to remove the FRT-flanked *kanR* cassette in MG1655 Δ*recA::kanR* and MG1655 Δ*lacI-Z::kanR* intermediate strains, respectively. All plasmids were constructed using standard cloning techniques with all information in Table S1.

### Genome targeting transformation assay in *E. coli*

Cells expressing Cas9 (**Figs. 1, 2, 4, S1, and S3**), eSpCas9(1.1) (**Fig. 2C**), SpCas9_HF1 (**Fig. 2C**), SpCas9_HeF (**Fig. 2C**), or AsCas12a (**Fig. 3, 5, S2, and S3**) were used to assess genome targeting in MG1655 and MG1655 Δ*recA*. The strains were struck to isolation on a Luria-Bertani (LB) plate with the appropriate antibiotics. Single colonies were inoculated into liquid LB with antibiotics for overnight culturing at 37°C shaking at 220rpm. Each colony represents a biological replicate. The following morning, cells were back-diluted into liquid LB with antibiotics and grown to an ABS_600_ of 0.6 - 0.8. Subsequently, the cells were made electrocompetent and transformed via electroporation with 50 ng of the appropriate gRNA plasmid (see Table S1 for more details). Cells were recovered in 500μL of Super Optimal broth with Catabolite repression (SOC) medium for one hour at 37°C shaking at 220rpm and then plated in five µL spot, serial dilutions or using full plate dilutions on LB agar with appropriate antibiotics. The cells were incubated for 16 hours at 37°C. The number of colonies were counted the following morning and the number of transformants was back-calculated.

### Editing assay in *E. coli*

To evaluate the editing efficiencies in *E. coli*, the same procedure was followed as in the transformation assay with small modifications. The cells were made competent at an ABS_600_ between 0.6 - 0.8 for experiments presented in Figs. 2B, 3C, 3D, and 5C. The cells were made competent at an ABS_600_ of ∼1.6 for experiments presented in Fig. 1D, 2C, 2E, and S2D. After transformation and selection on appropriate antibiotic plates, ten colonies from cells transformed with targeting gRNA plasmids and three colonies from cells transformed with the non-targeting gRNA plasmids were picked at random and re-struck onto LB plates with the appropriate antibiotics. These re-struck cells were incubated at 37°C for 16 hours. Colony PCRs were performed on the re-struck colonies depending on the target gene, PCR amplicons were then verified on a gel, and subject to PCR cleanup. The cleaned PCR products were then subjected to digestion by the appropriate restriction enzyme to evaluate if precise editing occurred. The digested products were resolved on agarose gels and the presence of a digestion banding pattern resulted in the colony being considered “edited”. No presence of the digestion banding pattern resulted in the colony being considered “non-edited”.

AvrII was used to evaluate precise editing for the RT used in Fig. 1D. BsgI was used to evaluate precise editing for the RT used in Figs. 2B, C, and E. AclI was used to evaluate precise editing for the RT used in the left panel of Fig. 3C. RE digests were set up according to the manufacturer’s protocol. Sanger sequencing was performed to evaluate precise editing for the RT used in the right panel of Fig. 3C when deleting 2nt. Colony PCR was used to evaluate precise editing for the RTs used in Fig. 3D when deleting the *lacZ* gene or replacing *lacZ* with *gfp*. A non-selective out-growth was used for the aTc inducible sgRNA editing experiment, Fig 2B. The transformation assay was performed as previously described through the recovery step in SOC. Upon recovery, 20μL of the recovering cells were back-diluted 100x into non-selective outgrowth medium (LB with chloramphenicol (cm, 34 μg/mL) and the appropriate amount of aTc) and incubated at 37°C shaking at 220rpm for 16 hours. The cells were then serially diluted in 1xPBS and 5μL aliquots of each dilution were plated on LB plates with cm and kanamycin (kan, 50 μg/mL) without further aTc induction.

### Editing assay in *K. oxytoca*

*Klebsiella oxytoca* MK01 was cultured in LB medium to ABS_600_ of 0.6 - 0.8. Cells were then made electrocompetent and transformed with 50 ng of pCas9KP plasmid. 50µL competent cells were recovered with 950µL liquid LB for one hour at 30°C shaking at 220 rpm. The culture was centrifuged at 8000xg for 3 minutes. After the supernatant was removed, cells were resuspended with 50µL LB and plated on LB plates supplemented with 60µg/mL apramycin. 3 single colonies representing biological replicates were inoculated in LB with apramycin and cultured at 30°C for 16 hours. 1mL of overnight cultures were inoculated into 100mL fresh LB and grown until ABS_600_ of 0.15 - 0.2, then induced with 10mL 20% L-arabinose LB supplemented with apramycin during which cultures were kept at room temperature shaking at 50 rpm for 2 hours (ABS_600_ of 0.6 - 0.8) and subsequently made electrocompetent as described before.

To evaluate editing efficiencies presented in Figure 4B, L-ara induced competent cells were co-transformed via electroporation with 200ng of sgRNA plasmids and ≈ 500ng of linear repair template assembled with 750-750bp of upstream and downstream homology arms adjacent to the gene-of-interest using SOE-PCR. Cells were recovered with 950µL LB for one hour at 30°C shaking at 220 rpm and then plated in serial dilutions on LB supplemented with apramycin (60µg/mL) and spectinomycin (300µg/mL). After overnight incubation at 30°C, 10 and 3 colonies were randomly picked from targeting sgRNA and non-targeting sgRNA containing transformation plates respectively and streaked out to fresh LB plates with the appropriate antibiotics and incubated at 30°C for 16 hours. The following day, colony PCRs were performed on the streaked-out colonies targeting the gene-of-interest. Successful gene deletion was verified on 1% TAE agarose gel run at 130V for 15 minutes and subsequently editing efficiencies were quantified.

### Editing assay in *K. pneumoniae*

The transformation assay was followed with small changes for improving transformation efficiency in *Klebsiella pneumoniae* ATCC 10031. Specifically, cells expressing the AsCas12a nuclease codon optimized for this bacterium have been back-diluted in LB medium with cm and 0.7 mM EDTA and made competent at ABS_600_ ≈ 0.4. The plasmid encoding the gRNA has been transformed as previously stated. For the editing experiment, a recombineering template with homology arms of 250 bp was cloned into the AsCas12a gRNA backbone prior to transformation. Ten and three colonies for the targeting and non-targeting samples respectively were picked for each biological replicate to be struck out on cm + kan agar plates for subsequent screening by colony PCR and Sanger sequencing. Some of these colonies were struck out on ampicillin (amp, 100 μg/mL) plates in parallel to check for resistance or sensitivity to amp respectively for unedited and edited colonies, Fig. 5D. One non-edited colony transformed with the NT gRNA plasmid and one edited colony obtained by using the mutR6 gRNA plasmid were made competent and transformed with pAmpR (i.e. CBS-3946) or with water and plated either on amp or LB only agar plates. Colony PCR was performed for three edited colonies transformed with pAmpR and three transformed with water. 10 µL of the purified PCR products were run on a 2% agarose TAE gel at 80V for 40 min and stained for 15 min in EtBr prior to visualization.

### Northern blotting analysis

MG1655 Δ*lacI-Z* cells were transformed with pCas9 and the appropriate gRNA plasmid (Table S1; pDC786, pDC841, pDC876, pDC869, pDC829, pDC860, pDC889, pDC862, and pDC886) and selected for on appropriate antibiotics. Individual colonies were inoculated into LB with appropriate antibiotics and incubated at 37°C shaking at 220rpm overnight. Cultures were then back-diluted into 15mLs of fresh LB with antibiotics and grown to an ABS_600_ of 0.6-0.8. 5mLs of this culture was mixed with 1mL of STOP mix, 95% of 100% ethanol and 5% hot phenol (Carl Roth Cat No. A980.1). The cultures were then snap frozen in liquid nitrogen and stored at -80°C until analysis. The frozen cultures were then thawed on ice for 1 hour. The cultures were spun down at 4700 rpm at 4°C for 20 minutes. The supernatant was removed and then subsequently spun down at 4700 rpm at 4°C for 1 minute. The supernatant was removed and the pellet was mixed with 600µL of lysozyme-solution (0.5mg/mL in TE buffer, pH 8.0). 60µL of 10% SDS was added and incubated at 64°C for 1 minute. 66µL of 1M sodium acetate (pH 5.2) was mixed into the solution and then 750µL of phenol (Carl Roth, Cat. no. A980.1) was added. The samples were incubated at 64°C and vortexed briefly every 30 seconds for 6 minutes. The samples were then incubated on ice for 3-5 minutes and spun down at 13,000 rpm at 4°C for 15 minutes. The upper phase was then transferred into the 2mL PLG tube (QuantaBio, Cat. no. 2302830). 750µL of chloroform (Carl Roth, Cat. no. 3313.1) was then added and incubated at room temperature for 3 minutes. The samples were then spun down at 13,000 rpm at 12°C for 15 minutes. The upper phase was then transferred into a sterile 2mL eppendorf tube where 1.4mL of 30:1 ethanol: sodium acetate pH 6.5 was added. The RNA was precipitated at -20°C for 2 hours. The samples were then spun down at 13,000rpm at 4°C for 30 minutes. The supernatant was removed and 500µL of 70% ethanol was added to the pellet. The samples were spun down at 13,000rpm at 4°C for 10 minutes. The supernatant was removed and pellets were allowed to air dry at room temperature with the lid open. The pellets were dissolved in 75-100µL nuclease-free water and allowed to incubate at 70°C shaking for 5 minutes. 10µg of each RNA sample were mixed with 2x GLII loading buffer (NEB, Cat. no. B0363A) loaded on an 8% polyacrylamide (Rotiphorese® Gel 40 (19:1) Acrylamide and Bisacrylamide solution Carl Roth, Cat. no.#3030.1) 7M Urea gel. The samples were run on the gel at 300V for 135 minutes using a gel transfer system (Doppel-Gelsystem Twin L, PerfectBlue).The samples were transferred to a Hybond-XL membrane (GE healthcare, RPN203S) at 50V for 1 hour at 4°C and then cross-linked using 0.12J (UV-lamp T8C; 254 nm, 8W). The membrane was hybridized overnight in 15mL of Roti-Hybrid Quick buffer (Carl Roth, Cat. no. A981.1) at 42°C with 2.5-10pmol/µL of γ 32P end-labeled oligodeoxyribonucleotides (Table S1), then wrapped into a clear foil and exposed in the cassette with the phosphor screen for 3 days and visualized on a Phosphorimager (Typhoon FLA 7000, GE Healthcare). prDC1416 was used to generate the blot in Fig. 1C, prDC1417 was used to generate Fig. S2C, and prDC1418 was used to generate Fig. S2D.

## Acknowledgments

The authors thank Atul K. Singh for generating the *E. coli* Δ*recA* strain used in this work. The pCas9 and pCRISPR plasmids were gifts from Luciano Marraffini (Addgene #: 42876 and 42875 respectively). The pCP20 plasmid was a gift from Jörg Vogel. The plasmid carrying the *ampR* gene used in the *K. pneumoniae* experiment was kindly cloned by Katharina G. Wandera and was sent to Addgene (# 184844) for a separate publication. **Funding**: This work was funded by the Joint Programming Initiative on Antimicrobial Resistance (01KI1824 to C.L.B. and T.S.).

## Author contributions

D.C., E.V., and C.L.B devised the study. D.C., E.V. J.Y., K.C., E. A., A.R., and T.A. performed experiments. D.C. and C.L.B wrote the manuscript with input from all other authors. E.V. generated the figures. T.S. and C.L.B acquired funding and supervised the project.

## SUPPLEMENTARY INFORMATION

### Supplementary figures

**Figure S1.**
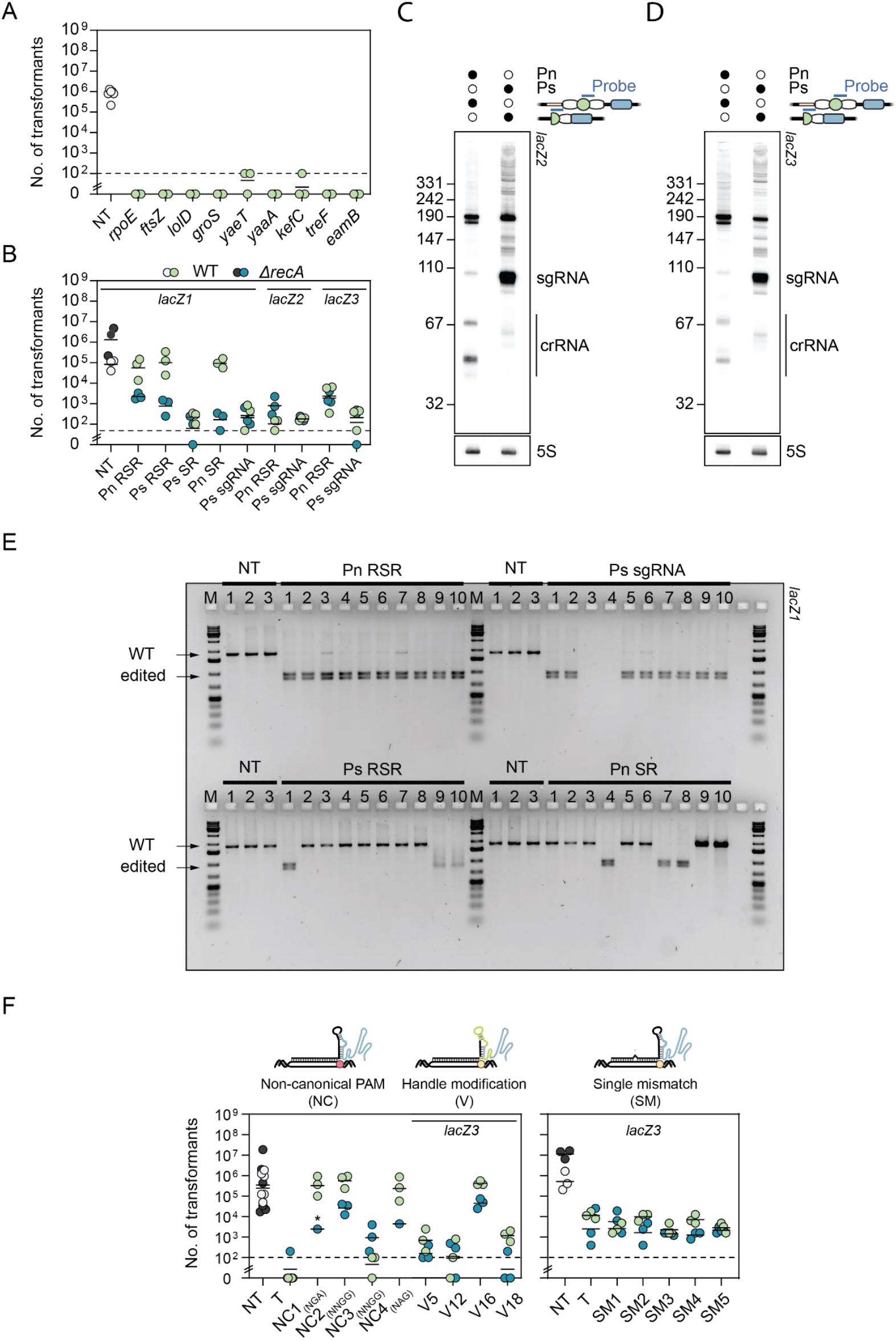
Assessing genome targeting and editing activity in *E. coli* with Cas9. **A)** Transformation assay in the WT strain of *E. coli* with Cas9 and sgRNAs. **B)** Transformation assays in the WT and Δ*recA* strain of *E. coli* with Cas9 and *lacZ* gRNA formats used for NBs. **C)** NB with *lacZ2* probe. **D)** NB with *lacZ3* probe. **E)** Gel readout of restriction enzyme digestion reactions on transformants with gRNAs and RTs from Figure 1D. **F)** Transformation assay in the WT and Δ*recA* strain of *E. coli* with Cas9 with modified gRNAs. * indicates that the transformants resulted in a lawn or uncountable colonies.

**Figure S2.**
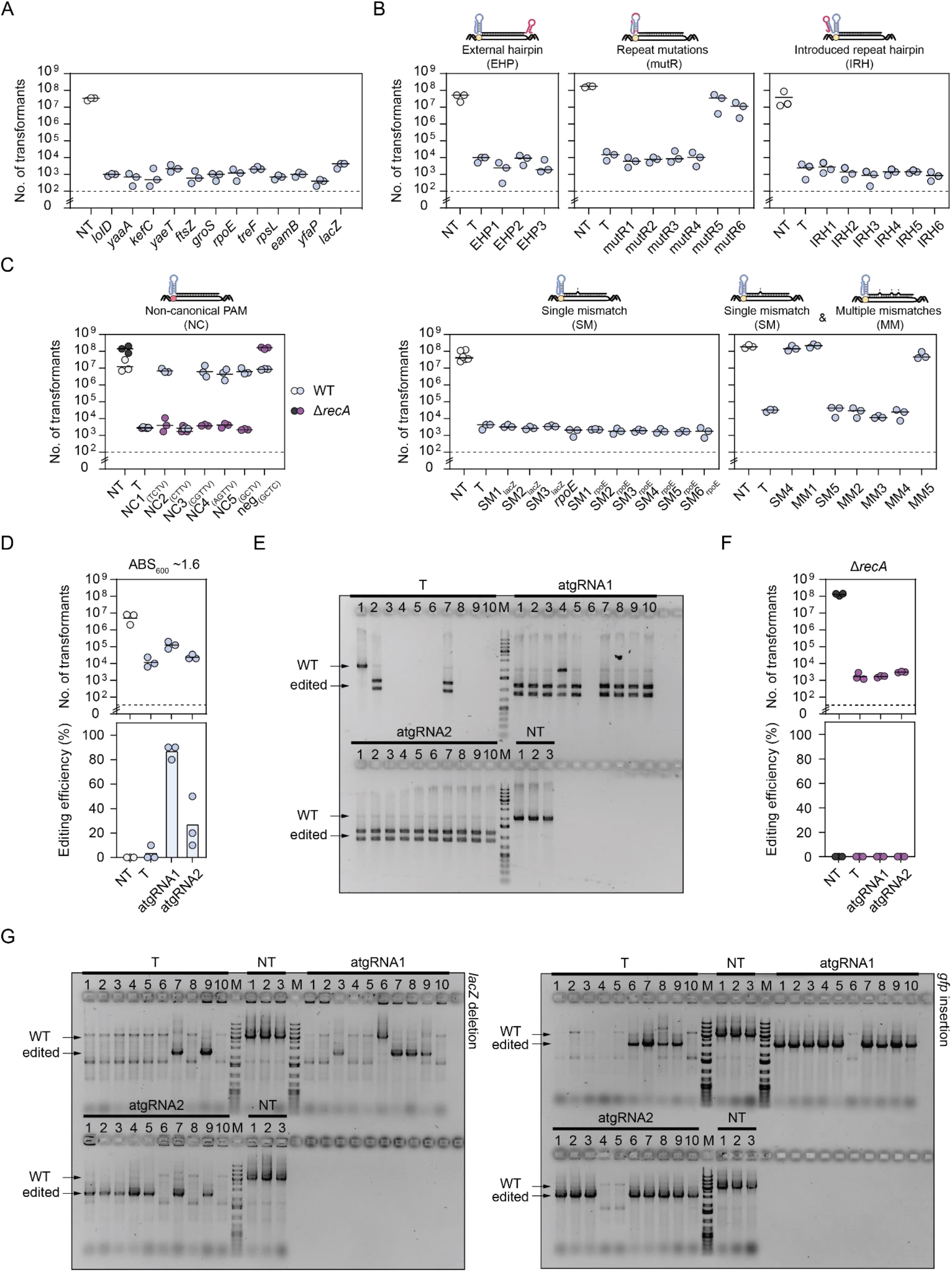
Assessing genome targeting activity in *E. coli* with Cas12a. **A)** Targeting in *E. coli* with AsCas12a. **B)** Transformation assay in WT strain of *E. coli* with AsCas12a using atgRNAs with an external hairpin, introduced repeat hairpin, repeat mutations, and single and multiple mismatches in the *lacZ* guide. Single mismatches were also generated for a guide targeting the *rpoE* gene. **C)** Transformation assay in WT and Δ*recA* strain of *E. coli* with AsCas12a and less-preferred PAMs. neg represents a target site adjacent to a non-PAM sequence. **D)** Editing assay to introduce an AclI restriction enzyme site into *lacZ* at ABS_600_ ≈ 1.6. The experiment showed that for AsCas12a editing cells in the stationary phase led to less transformants for the atgRNAs and lower editing efficiency compared to the experiment performed in exponential phase in Fig 3C. Each dot in the lower graph represents the fraction of edited cells for 10 and 3 screened colonies for the targeting and NT samples respectively. **E)** Representative gel for screening the insertion of an AcII restriction site in the *lacZ* gene. Related to Figure 3C. **F)** Editing assay to an AclI restriction enzyme site into *lacZ* in the *E. coli* Δ*recA* strain. Each dot in the editing efficiency graph represents the value obtained for 3 colonies (n=3). **G)** Gel readouts for *lacZ* deletion and gfp insertion with 750 bp homology arms.

**Figure S3.**
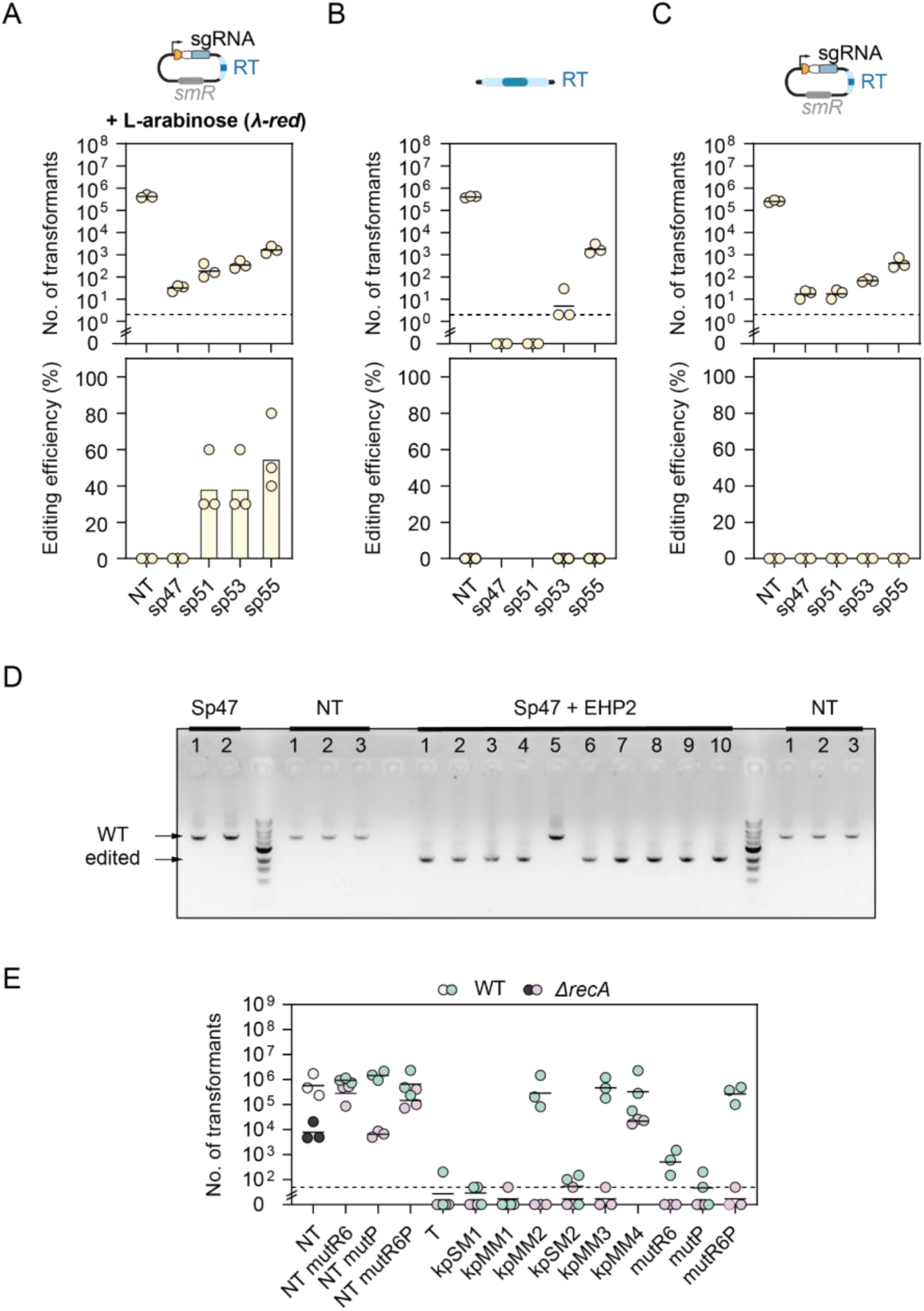
Assessing genome targeting and editing activity in strains of Klebsiella. **A)** Editing assay in the WT *K. oxytoca* with a plasmid-based RT and lambda-red induction. **B)** Editing assay in the WT *K. oxytoca* with linear RT and no lambda-RED induction. **C)** Editing assay in the WT *K. oxytoca* w/plasmid-based RT and no lambda-RED induction. **D)** Gel image of colony PCR to screen for *cydAB* deletion from Fig. 4B. **E)** Targeting assay in *K. pneumoniae* with different guides without the RT. mutP indicates a point mutation in the constitutive promoter, while mutR6P indicates both the mutation of the repeat mutR6 and the promoter mutation mutP.

### List of SI tables

Table S1. Strain, plasmids, and oligos.

